# Pre-synaptic inhibition of afferent feedback in the macaque spinal cord does not modulate with cycles of peripheral oscillations around 10 Hz

**DOI:** 10.1101/012583

**Authors:** Ferran Galán, Stuart N Baker

**Affiliations:** Institute of Neuroscience, Newcastle University, Newcastle upon Tyne NE2 4HH, UK

**Keywords:** primary afferent depolarization, pre-synaptic inhibition, tremor

## Abstract

Spinal interneurons are partially phase-locked to physiological tremor around 10Hz. The phase of spinal interneuron activity is approximately opposite to descending drive to motoneurons, leading to partial phase cancellation and tremor reduction. Pre-synaptic inhibition of afferent feedback modulates during voluntary movements, but it is not known whether it tracks more rapid fluctuations in motor output such as during tremor.

In this study, dorsal root potentials (DRPs) were recorded from the C8 and T1 roots in two macaque monkeys following intra-spinal micro-stimulation (random inter-stimulus interval 1.5-2.5 s, 30-100μA), whilst the animals performed an index finger flexion task which elicited peripheral oscillations around 10Hz. Forty one responses were identified with latency <5ms; these were narrow (mean width 0.59 ms), and likely resulted from antidromic activation of afferents following stimulation near terminals. Significant modulation during task performance occurred in 16/41 responses, reflecting terminal excitability changes generated by pre-synaptic inhibition (Wall’s excitability test). Stimuli falling during large-amplitude 8-12Hz oscillations in finger acceleration were extracted, and sub-averages of DRPs constructed for stimuli delivered at different oscillation phases. Although some apparent phase-dependent modulation was seen, this was not above the level expected by chance.

We conclude that although terminal excitability reflecting pre-synaptic inhibition of afferents modulates over the timescale of a voluntary movement, it does not follow more rapid changes in motor output. This suggests that pre-synaptic inhibition is not part of the spinal systems for tremor reduction described previously, and that it plays a role in overall – but not moment-by-moment – regulation of feedback gain.

## Introduction

Physiological tremor is produced by multiple interacting mechanisms. These include mechanical resonance of limb articulations (Marsden et al. 1969), and oscillations in the stretch reflex loop consequent on the peripheral conduction delays (Lippold 1970). However, there is also a centrally generated component in the 8-12Hz frequency range, as its frequency is unaltered by manoeuvers such as loading the limb, which alter mechanical and reflex resonant frequencies (Elble and Randall 1978). Studies assessing slow finger movements in non-human primates have found coherence at ∼10Hz between acceleration and the activity of multiple motor structures during both active movements and periods of steady holding (Williams et al. 2010a), suggesting that physiological tremor and discontinuities during slow finger movements reflect the same underlying phenomenon. Interestingly, motor cortical oscillations at ∼10Hz are not coherent with muscle activity in this range during steady holding (Baker et al. 1997; Conway et al. 1995; Salenius et al. 1997) despite their passage down the corticospinal tract (Baker et al. 2003). These observations have suggested the existence of an active neural filter, which removes ∼10Hz components from the input to motoneurons (Williams and Baker 2009; Williams et al. 2010a), that could be important in the reduction of tremor.

Spinal networks could influence motoneurons by multiple possible pathways, most obviously by excitatory or inhibitory synapses on the motoneurons themselves. One known instance of such a direct synaptic effect is Renshaw cell recurrent inhibition, which previous work has shown can partially cancel ∼10Hz components in motoneuron input (Williams and Baker 2009). However, spontaneous oscillations in the cord are also synchronized with similar oscillations in dorsal root potentials (Lidierth and Wall 1996), and are associated with primary afferent depolarization (PAD) (Manjarrez et al. 2000) which reflects pre-synaptic inhibition of afferent input. It is not known whether these spontaneous spinal oscillations in anesthetised animals are related to tremor circuits. Furthermore, it is known that muscle spindle afferents modulate their discharge with the phases of peripheral oscillations around 10 Hz (Baker et al. 2006; Wessberg and Vallbo 1995), and that pre-synaptic inhibition modulates during voluntary movements (Hultborn et al. 1987; Seki et al. 2003), suppressing motor oscillations during forelimb movement (Fink et al. 2014). Therefore, the modulation of presynaptic inhibition with the ∼10 Hz oscillations of tremor appears as a reasonable hypothesis.

Using a new technique that allowed recordings from a mixed population of muscle and cutaneous afferents in awake behaving primates, this study investigated whether afferent axon terminal excitability modulates during performance of a slow index finger flexion task, and with the cycles of physiological tremor which are prominent in such a task. Although robust modulation over the second-to-second timescale of task performance was regularly seen, the data contained no evidence for faster modulation during the tremor cycle. These results suggest that pre-synaptic inhibition may act as a less temporally-precise gate for afferent inflow, but does not sculpt sensory input and its reflex consequences over timescales comparable to endogenous oscillations in motor output.

## Methods

### Behavioural task

Two female *Macaca mulatta* monkeys (denoted I and V) were trained to perform a finger flexion task for food reward, similar to that used in previous work (Soteropoulos et al. 2012; Williams et al. 2009; Williams et al. 2010a). The index finger of the right hand was inserted into a narrow tube, which restricted movement to the metacarpophalangeal (MCP) joint. The tube was attached to a lever that rotated coaxially with the MCP joint; a motor exerted torque in a direction to oppose flexion. Lever angular displacement was sensed by an optical encoder and fed back to the animal via a cursor on a computer screen. A displacement of 0° indicated the neutral position, where the finger was in the same plane as the palm. Positive angles denoted finger flexion. During each trial the palm and digits 1, 3, 4, and 5 lay horizontally against a flat surface and the elbow and upper arm were held in a sleeve. The contralateral arm was unrestrained, and remained at rest during task performance; at the end of a successful trial, the contralateral arm retrieved the food reward.

For both animals, a trial commenced when a rectangular target appeared at 8° displacement. The animal moved the cursor into this target, which then moved over a linearly increasing displacement (ramp). In monkey I, the ramp phase lasted 1.5 s, with final displacement 20°; the trial was completed at the end of the ramp phase. In monkey V, the ramp phase lasted 2 s, with final displacement 16°; at the end of the ramp, there was a hold phase of constant target displacement lasting 1 s. Maintenance of the cursor within the target (allowed error ±1.4°) led to a food reward. An accelerometer attached to the lever measured movement discontinuities during the target ramp (band-pass, 1–100 Hz).

### Surgical preparation

Following training, both animals were implanted under general anaesthesia (3.0–5.0% sevoflurane inhalation, intravenous infusion of 0.025 mg.kg^-1^.h^-1^ alfentanil) and aseptic conditions with a stainless steel headpiece for head fixation. After an appropriate recovery period, a further surgery implanted a spinal chamber over a laminectomy spanning vertebrae C5-C7, together with a bipolar cuff electrode on the C8 (monkey I) or T1 (monkey V) dorsal root adjacent to the cord. This cylindrical cuff electrode was modelled on those commonly used for peripheral nerve stimulation and recording. It was manufactured from flexible medical-grade silicone polymer to have an internal diameter of 2.0 mm and length 5 mm. The cuff contained two platinum electrodes which ran around the internal circumference (electrode width 1.0 mm, separation 1.5 mm), and were spot-welded to Teflon-insulated stainless steel wire (wire diameter 150 μm). The dorsal root was inserted into the cuff via a slit along the length of the cuff, which was then closed using two silk sutures which ran around the outside. Cuff placement was made possible by the fact that in monkey the C8/T1 roots run parallel to the cord for some distance before turning to exit the facet joint. This displacement between spinal segment and equivalent vertebra is much more marked in monkey than in man, where roots do not run parallel to the cord in this way until the mid-throracic level. The wires were run over the lateral mass and then up the side of the chamber, where they terminated in a connector. Both wires and connector were covered in dental acrylic for protection. Several features of the cuff design were intended to record selectively from the dorsal root, whilst reducing potentials from the adjacent cord. Firstly, the electrodes were closely spaced within an insulating cylinder, with the distance from each electrode to the cuff edge similar to the inter-electrode spacing. Secondly, the cuff lay parallel to the cord, so that each electrode was equidistant from any cord generators. We would therefore expect that any residual potentials from the cord would be similar on each electrode, and hence cancel in the differential recording.

A full program of post-operative analgesia followed all surgical procedures; all procedures were performed under appropriate licences issued by the UK Home Office under the Animals (Scientific Procedures) Act (1986) and were approved by the Animal Welfare and Ethical Review Board of Newcastle University.

### Recordings

During task performance both head and spinal implants were fixed to the primate chair and a microdrive was interfaced to the spinal chamber via an X-Y positioning stage. Differential recordings from the contacts of the dorsal root cuff electrodes yielded the dorsal root potential (DRP, band pass 3 Hz-2 kHz, gain 50 K). Intraspinal microstimulation (ISMS) (bipolar pulses, 0.07-0.1 ms per phase, inter-stimulus interval chosen at random from a uniform distribution in the range 1.5-2.5 s, 30-100 μA) was delivered through a tungsten microelectrode (impedance 1 MΩ) inserted into the spinal grey matter at a location which evoked responses in the DRP (see Fig. 1). Electrode depth ranged 2.0-7.2 mm relative to the surface of the dura mater (mean 5.1 mm, SD 1.2 mm). Stimulus intensity was set to yield response amplitudes around half of the maximum. Stimulus timing was controlled by a 1401 interface (CED Ltd, Cambridge, UK), which also recorded DRP (sampling rate 20 kHz), lever position and acceleration (sampling rate 1 kHz) and task markers to disk for later off-line analysis.

**Figure 1:**
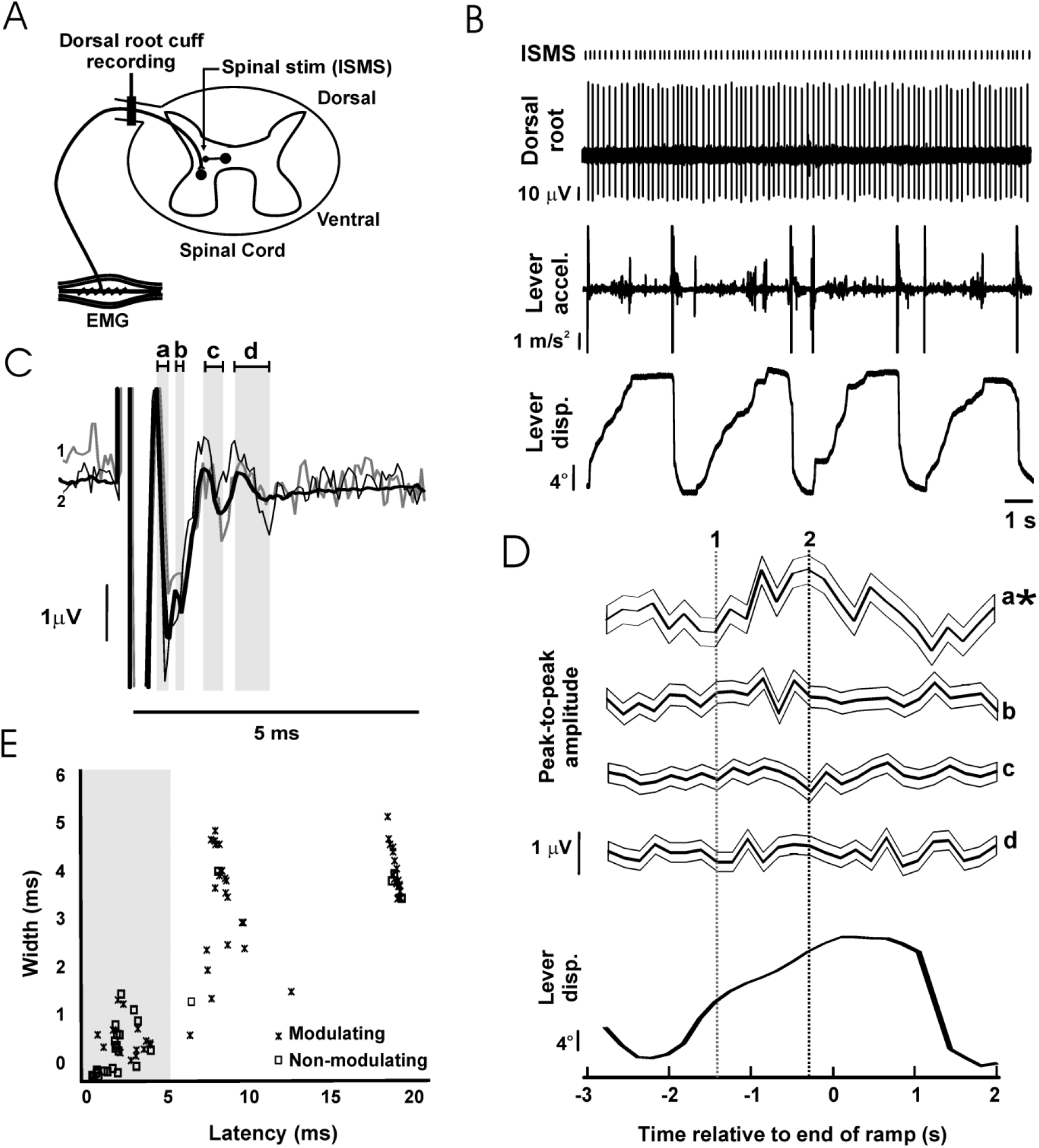
Dorsal root potentials evoked by intra-spinal micro-stimulation (ISMS). A, schematic of the recording setup. A bipolar cuff electrode was implanted on the C8 or T1 dorsal root adjacent to the spinal cord. During task performance, ISMS was delivered at a location which elicited antidromic responses in the mixed population of afferents recorded at the dorsal root electrode. B, example raw data during task performance. C, example of responses evoked at the dorsal root by ISMS to a single spinal site (65 μA). Thick black trace represents grand-average (n=4698 stimuli); thin black and thin grey traces represent sub-averages from task-dependent bins marked by dotted lines in (D). Grey shading and lower case letters indicate different response components. D, task-dependent modulation of the responses indicated by lower case letters in (C). Each trace shows the mean response, with faint surrounding traces indicating the SEM. Traces are aligned to the end of the ramp phase of the task; average lever displacement is shown below in the same timeframe for comparison. Asterisks denote responses with significant task-dependent modulation. Vertical lines indicate times used to compute sub-averages illustrated in (C). E, scatter plot of the width versus latency of responses (n=88). Only responses with latency shorter than 5 ms (grey shading) were used in subsequent analysis (n=41).

Dorsal root responses following ISMS had a complex profile. In order to clarify the origin of the different components, these were compared with responses following median nerve stimulation (1ms bipolar pulses, 1-3 Hz, 0.4-1.8 mA applied to surface electrodes at the wrist) in both the dorsal root cuff and the spinal cord (recorded by a tungsten microelectrode as above, band pass 1 Hz-5 kHz, gain 2 K, sampling rate 10 kHz; Fig. 2).

**Figure 2:**
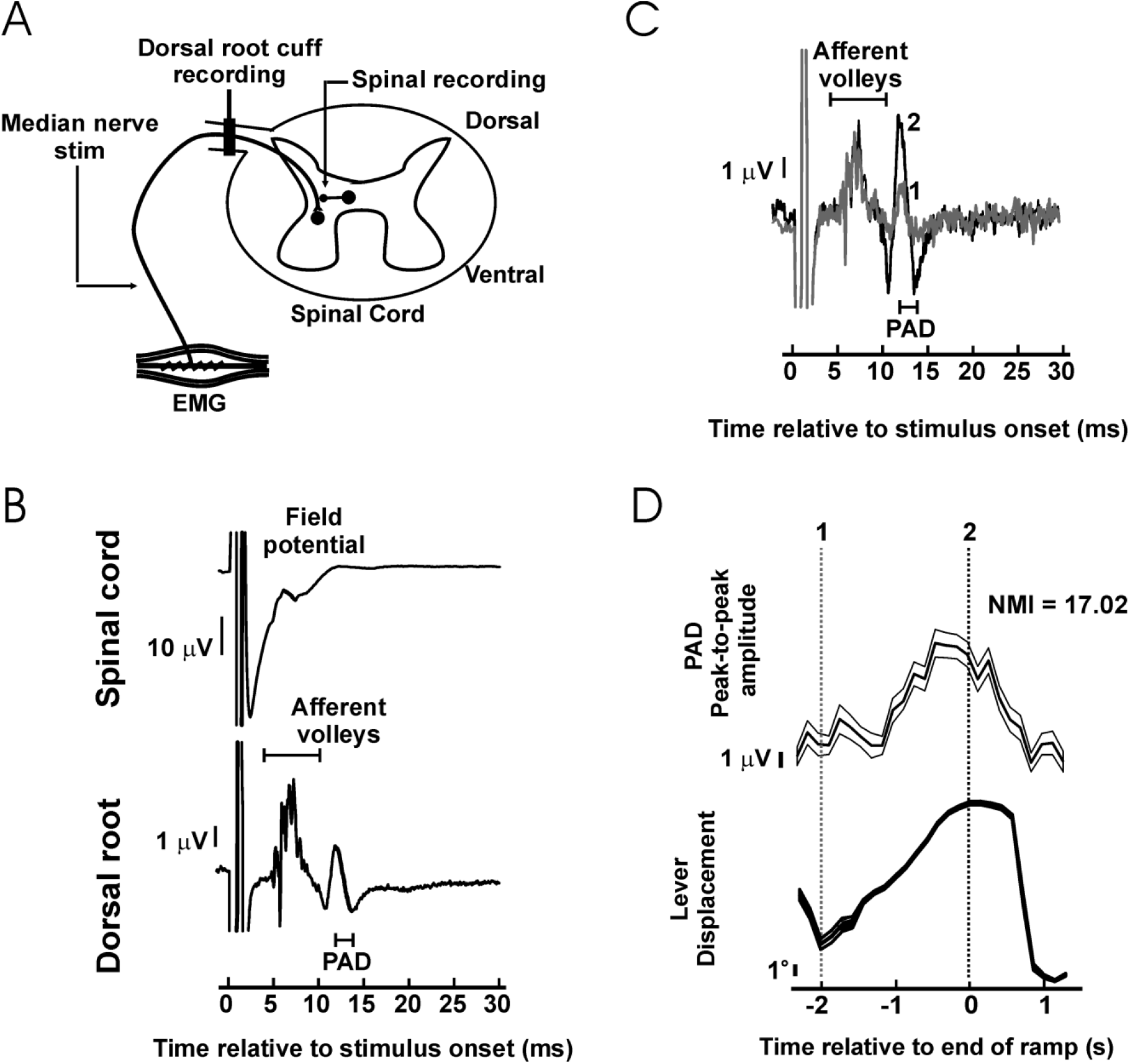
Dorsal root potentials evoked by peripheral nerve stimulation. A, during task performance the median nerve was stimulated whilst recordings were made from the dorsal root and the spinal cord. B, example of averaged responses in one experiment. In the dorsal root potential, an early afferent volley is followed by later primary afferent depolarisation potential (PAD). In the spinal cord, the field potential developed at latencies comprised between the early afferent volleys and PAD in dorsal root. Thick black trace represent grand-average (n=10859 stimuli). C, dorsal root potentials during task performance, black and grey traces represent sub-averages from the times indicated in (D). D, significant task-dependent modulation of the amplitude of the PAD potential shown in (B-C). Top trace is mean PAD amplitude (thick line) and its SEM (thin line). Bottom trace is the average lever displacement, in the same timeframe, for reference. Vertical dotted lines indicate time points used for the corresponding coloured sub-averages in (C).

### Modulation of ISMS evoked responses at dorsal root

Evoked responses in the DRP following ISMS were first determined by averaging triggered by all stimuli delivered to a given spinal site. This allowed estimation of the peak-to-peak amplitude of each component, measured over a time window selected manually. The latency was estimated from the time of the first peak of that component.

To estimate the task-dependent modulation of responses, trials were first aligned to the end of the ramp phase of the task. Stimuli were then sorted depending on when they had occurred relative to this alignment point, into 26 non-overlapping bins, each 200 ms wide, spanning from 3 s before to 2 s after the end of the ramp. Selective averages were compiled of stimuli in each bin, and amplitude measured from each average using the time window defined from the all-stimulus average.

To estimate how evoked responses modulated with tremor phase, stimuli were first selected using two criteria. It is known that peripheral oscillations are stronger during periods of finger movement (which motivated our use of a task incorporating a ramp phase). Accordingly, the finger lever velocity *V* was estimated over a window prior to the stimulus at time *T* as

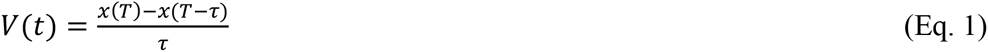

Where *x* represents the lever displacement, and the window length *τ* was set to 100 ms. The amplitude spectrum of the lever acceleration was estimated over a window prior to the stimulus at time *t* (where *t*<0) using non-symmetric causal wavelets as described in (Mitchell et al. 2007). In brief, the wavelet *W* at frequency *f* was defined as the product of an alpha function with peak 0.8/*f* before the stimulus, with a complex sinusoid:

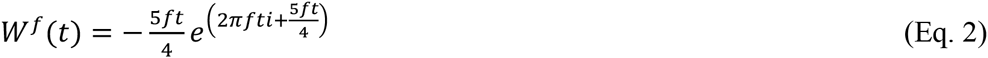

For a given frequency *f*, a section of accelerometer signal *S* was extracted lasting seven oscillation periods prior to the stimulus at time *T*. The dot-product of the accelerometer signal *S* with the wavelet *W*^*f*^ was found:

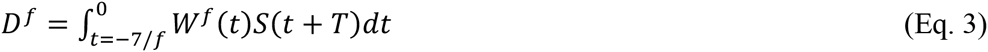

The amplitude A and phase φ were measured as:

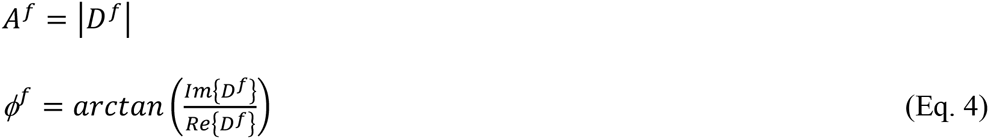

Only stimuli with V>15 °/s and which had the bin with largest spectral power lying within the 8-12 Hz range were included for subsequent analysis of tremor modulation. Note that using these criteria, stimuli falling in unsuccessful trials (e.g. where the animal strayed outside the imposed limits on tracking performance towards the end of a trial) were able to be used as well as those during successful task performance.

Stimuli which survived this pre-selection were then grouped by the phase of on-going lever acceleration oscillations in which they occurred, using eight equally-sized bins from 0 to 2π. DRP averages were then compiled selectively from stimuli in each bin, and response amplitude measured for each sub-average as for the determination of task modulation. Because the wavelet analysis of Eq. 3 used only acceleration data before the stimulus, any consequence of the stimulus on the periphery (such as a twitch) could not affect the phase determination. Inter-stimulus intervals were chosen at random (range 1.5-2.5s) to prevent phase-locking of oscillations to the stimulus.

Plots of response amplitude versus bin number often showed complex patterns of modulation, for both task and phase dependent modulation. A simple summary measure, which quantified the overall extent of this modulation in a single number, was developed. First, a raw modulation index from the experimental data, *I*_*exp*_, was computed as

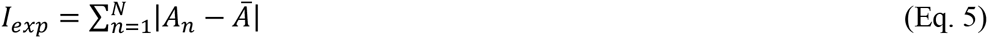

Where *A*_*n*_ is the response amplitude measured in bin *n*. The number of bins *N* was 26 for task modulation and 8 for tremor. The mean response *Ā* was calculated as:

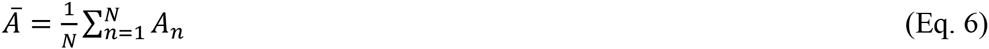

The measure *I* quantifies how much single bin responses deviate from the average response, but it is difficult to interpret the scale of this number. Accordingly, surrogate datasets were generated, which estimated how great *I* would be, on the null hypothesis of no response modulation above that expected by chance fluctuations. Surrogates were compiled by randomly shuffling bin assignments of individual stimuli; for a given bin, the number of stimuli assigned to it was fixed equal to the number in the experimentally determined dataset. Sub-averages were compiled for the surrogate data, and the measure *I* recompiled. This was repeated 500 times, using different random assignments of stimuli to bins. The mean *Ī* and standard deviation σ_*I*_ of the surrogate values of *I* was found, allowing us to compute a normalised modulation index (*NMI*_*exp*_) as:

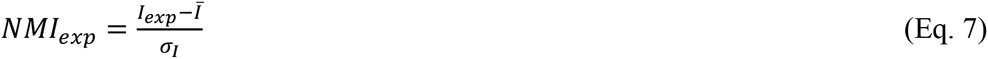

If *I*_*exp*_ exceeded the 95^th^ percentile of the surrogate values of *I*, the modulation was considered to be statistically significant (P<0.05).

To interpret the scale of tremor modulation in the 8-12 Hz range further and be able to compare it with other frequency ranges, data was combined across recording sites in two ways, estimating both the count *C* of significantly tremor-modulating responses, and also the mean modulation index 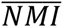 across all sites. Surrogate measures of C and 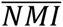 were generated by counting or averaging over one surrogate value of *I* per site; this was repeated 500 times with different randomly generated surrogates. If *C*_*exp*_ and 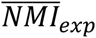 from the experimental data exceeded the 95^th^ percentile of the surrogate values, they were considered to be statistically significant (P<0.05). This whole procedure was repeated for frequencies between 6 and 50 Hz (1 Hz resolution).

### Identification of task-dependent modulation patterns of ISMS evoked responses at dorsal root

It is of interest to determine whether responses with a significant task-dependent modulation from different spinal sites could be grouped into a smaller number of representative profiles. Profiles were accordingly subjected to unsupervised *k-*means clustering (Jain 2010), using the correlation between profiles of amplitude versus bin number as the metric of pairwise distance. The number of identified patterns (=number of clusters, s) was chosen by maximizing the relatedness of the modulating responses across solutions for s=1…10. Response relatedness was estimated using the intra-cluster correlation coefficient:

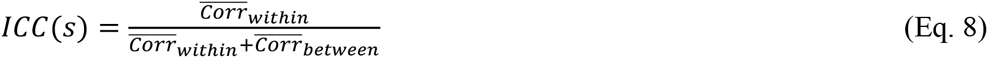

Where 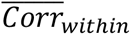 is the average squared correlation within clusters, and 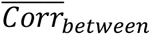 is the average squared correlation between clusters. Given a clustering solution *s* with clusters indexed by *c* or *d*=1..*s*, and each cluster containing *n*_*c*_ responses 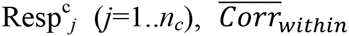 and 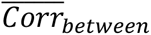 were calculated as:

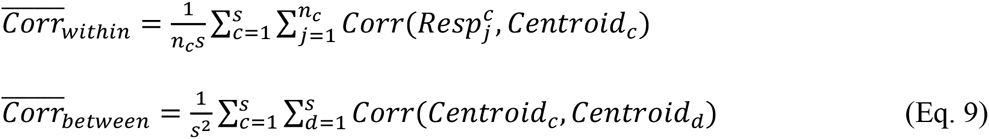

Where 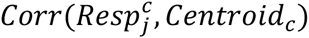 is the squared correlation between a response 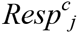 and the centroid of parent cluster, and *Corr*(*Centroid*_*c*_, *Centroid*_*d*_) is the squared correlation between centroids of clusters *c* and *d*.

Values of *ICC*_*s*_ close to one reflect solutions where responses are very similar within a cluster, but unrelated to those in another cluster. Estimates of *ICC*_*S*_ were produced using leave-one-out cross-validation, in which each response was correlated with cluster centroids determined after excluding that response from the dataset.

All analysis routines were implemented in the MATLAB package (The MathWorks Inc, Natick, MA, USA).

## Results

### Responses evoked in Dorsal Root Recordings by ISMS

Results were available from stimulation at a total of 21 spinal sites (8 monkey I, 13 monkey V). At each site, between 392 and 29516 stimuli were delivered (mean stimulus intensity 82 μA, SD 64 μA), whilst the animal performed between 20 and 161 successful trials of the task. Figure 1B illustrates typical raw data from an experiment, and Fig. 1C shows the averaged DRP evoked by the ISMS (stimulus intensity 65 μA, depth 4.4 mm). A complex waveform was visible in this average, reflecting multiple components of the response which are identified by the grey shading labelled with lower case letters.

Figure 1D shows how the different parts of the response from Fig. 1C modulated with task performance. Each trace shows the amplitude of one component as a function of time during the task (see averaged lever displacement beneath as a reference); traces illustrate both averaged amplitude (thick lines) and the corresponding standard error of the mean (thin lines). The earliest response (a) exhibited significant modulation with task, and the corresponding NMI value was above the 95^th^ percentile of those in surrogate datasets (experimental / 95^th^ centile surrogate NMI: 2.0/1.6). By contrast, later responses (b, c, d) had NMI below those expected by chance from surrogate data (b, 1.2/1.6;c, 1.3/1.8; d, 0.5/1.9), indicating no significant modulation with task. Across all 21 spinal sites which were stimulated, a total of 88 distinct responses were identified in the DRP, 57 of which (65%) modulated significantly with the task. In order to provide some insight into the physiological mechanisms generating the different response components, Fig. 1E plots their width (peak-trough time) versus latency (time of earliest peak/trough); components which modulated significantly with task are identified by crosses. It is clear that there are three broad classes of response. The earliest components (latency <5 ms; mean 2.44 ms, SD 1.15 ms) were narrow (width 0.59±0.45 ms, mean ± SD), and contained a mixture of modulating (16/41) and non-modulating (25/41) effects. The narrow nature of these responses suggests that these are most likely to reflect antidromic action potentials generated by direct stimulation of afferent axons within the cord. Such effects could exhibit a task relationship if the stimulating electrode was close to axon terminals, and those terminals were depolarized by axo-axonic synapses mediating primary afferent depolarization (PAD; Wall 1958), thereby modulating their excitability to the stimulus.

There appeared to be two later clusters of responses, with mean latencies of 8.1 ms and 18.1 ms. These showed a greater incidence of task-dependent modulation (22/24 and 19/23 responses respectively); both groupings of response were broader (widths 3.3±0.1 ms and 4.1±0.1 ms respectively). One possible cause for these effects could be PAD elicited in afferent axon terminals following activation of spinal neurons by the stimulus (either directly, or trans-synaptically), and passively conducted to the dorsal root recording site (Wall 1958). For the latest responses, it is possible that they are caused by reafference following a peripheral twitch induced by the ISMS.

### Comparison with Responses evoked in Dorsal Root Recordings by Peripheral Nerve Stimulation

Further insight into the mechanisms generating the later DRPs described above was provided by examining the responses to peripheral nerve stimulation (Fig. 2A). Figure 2B shows the average response after stimulation of the median nerve at the wrist, at an intensity above motor threshold that did not interfere with task performance (1.4 mA), for a single recording session. The DRP recording showed a compound volley, which has a first peak latency 3.9 ms after the stimulus. This contained multiple sharp components, presumably reflecting axons of different conduction velocities, and was followed by a slower potential (width peak-to-peak 1.0 ms), with first peak at 10.1 ms after the stimulus. We consider that this component is likely to reflect PAD (see Discussion). In the spinal cord, field potential onset was measured 4.2 ms after the stimulus. Figure 2C-D illustrates that this later potential modulated strongly with task performance, indicating that the excitability of neural populations mediating PAD changed in a task-dependent manner. Significant task-dependent modulation of a potential similar to that seen in Fig. 2B was observed in 3/4 recording sessions where median nerve stimulation was tested (NMI values 12.3, 5.0 and 17.0; probability of 3 or more out of 4 significant values by chance is P<5×10^-5^, binomial distribution).

In these three sessions, the average latency difference between the earliest afferent volley and the later PAD potential was 5.1±0.58 ms; the width of the later PAD potential was 1.1±0.05 ms (both mean±SEM). The latency is comparable to the later potentials seen following ISMS in Fig. 1E, although those elicited by afferent input were considerably narrower. It is possible that changes in terminal membrane conductances following prior activation by the afferent volley interacted with the PAD to reduce its width. The responses to median nerve stimulation therefore seem broadly to support the idea that the later responses to ISMS reflect PAD elicited by activation of spinal neurons.

### Patterns in the Task-dependent Modulation of ISMS-Evoked Dorsal Root Responses

A k-means clustering approach was used to examine whether there were any repeatable patterns in the task modulation profiles of the short-latency (<5 ms) ISMS responses from different sites (see Methods). A plot of the intra-cluster correlation (Fig. 3A) revealed a sharp increase in going from one to two clusters, but then only a small increase as the cluster number was further increased, peaking at four clusters. Figure 3B presents information on the different profiles identified. Of the 16 responses with significant modulation, 7 showed response facilitation during the task ramp phase (MP+), while 5 showed a facilitation just after the ramp phase ended (MP-). The remaining two clusters appeared to have erratic profiles and were categorized as MP*1 and MP*2 (n=2 sites each). Averaged lever displacement traces are shown at the bottom in Figure 3B, and make clear that there were differences in the temporal profile of task performance between the two animals. Interestingly, the modulation profiles MP+, MP*1 and MP*2 all occurred in monkey V, whereas MP- profiles all occurred in monkey I, suggesting that individual differences in the task and its performance lay behind the modulation differences. There was no significant difference between the modulation depths of the four patterns (Fig. 3C; P=0.104, Kruskal-Wallis test).

**Figure 3:**
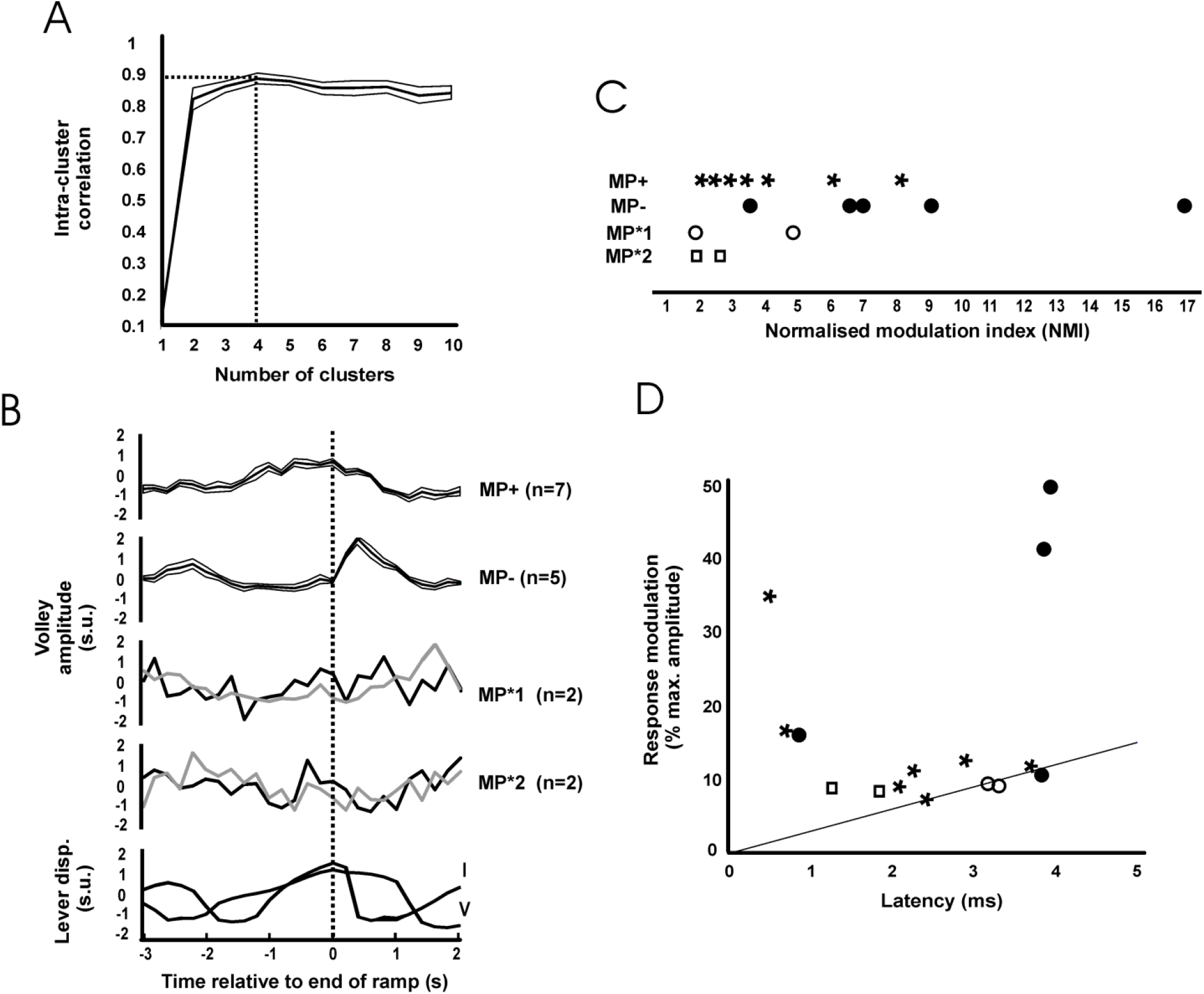
Clustering the patterns of task-dependent modulation. A, intra-cluster correlation (ICC, shown as mean ± SEM) as a function of the number of clusters. Maximum ICC was with 4 cluster (dotted lines). B, mean ± SEM (thick lines and associated thin lines) of MP+ and MP- and single responses of MP*1 and MP*2 modulating patterns identified from all 16 significantly modulating responses. Traces are aligned relative to the end of the ramp phase of the task. Beneath are shown averaged lever displacement traces for each monkey in the same timeframe, for comparison. In monkey I the ramp lasted 1 s, in monkey V it lasted 2 s. All traces have been normalised to have zero mean and unit standard deviation (standard units). C, normalised modulation index of significantly modulating responses, separated by cluster class. D, scatter plot of the modulation, expressed as the difference between the minimum and maximum response as a percentage of the maximum, versus response latency, for significantly modulating responses. The line represents the relationship expected from collision with orthodromic spikes modulating by 30 discharges per second (see Discussion).

Figure 3D presents the relation of the size of the modulation in response (calculated as the difference between minimum and maximum response, as a percentage of the maximum) with response latency. The straight line indicates the largest modulation which we estimate could be generated by collision between orthodromic and antidromic spikes; more detail on the basis for the calculation of this line and the implications of this plot are given in the Discussion.

### Modulation of ISMS Evoked Dorsal Root Responses with Tremor Cycle

Figure 4A illustrates typical raw data from an experiment, marking with vertical dotted lines the stimuli which were included for analysis of tremor modulation based on a linearly increasing lever displacement and a power spectral peak of lever acceleration in the 8-12 Hz range. Figure 4B shows the asymmetric wavelet used to extract amplitude and phase information from the acceleration signal (see Methods). Figure 4C shows two example phase-dependent modulation profiles of responses classified as MP+ and MP- on the basis of task. Each of these had NMI values above those expected by chance from surrogate data, indicating a significant modulation with tremor oscillatory phase at the frequencies illustrated (9 Hz and 10 Hz respectively).

**Figure 4:**
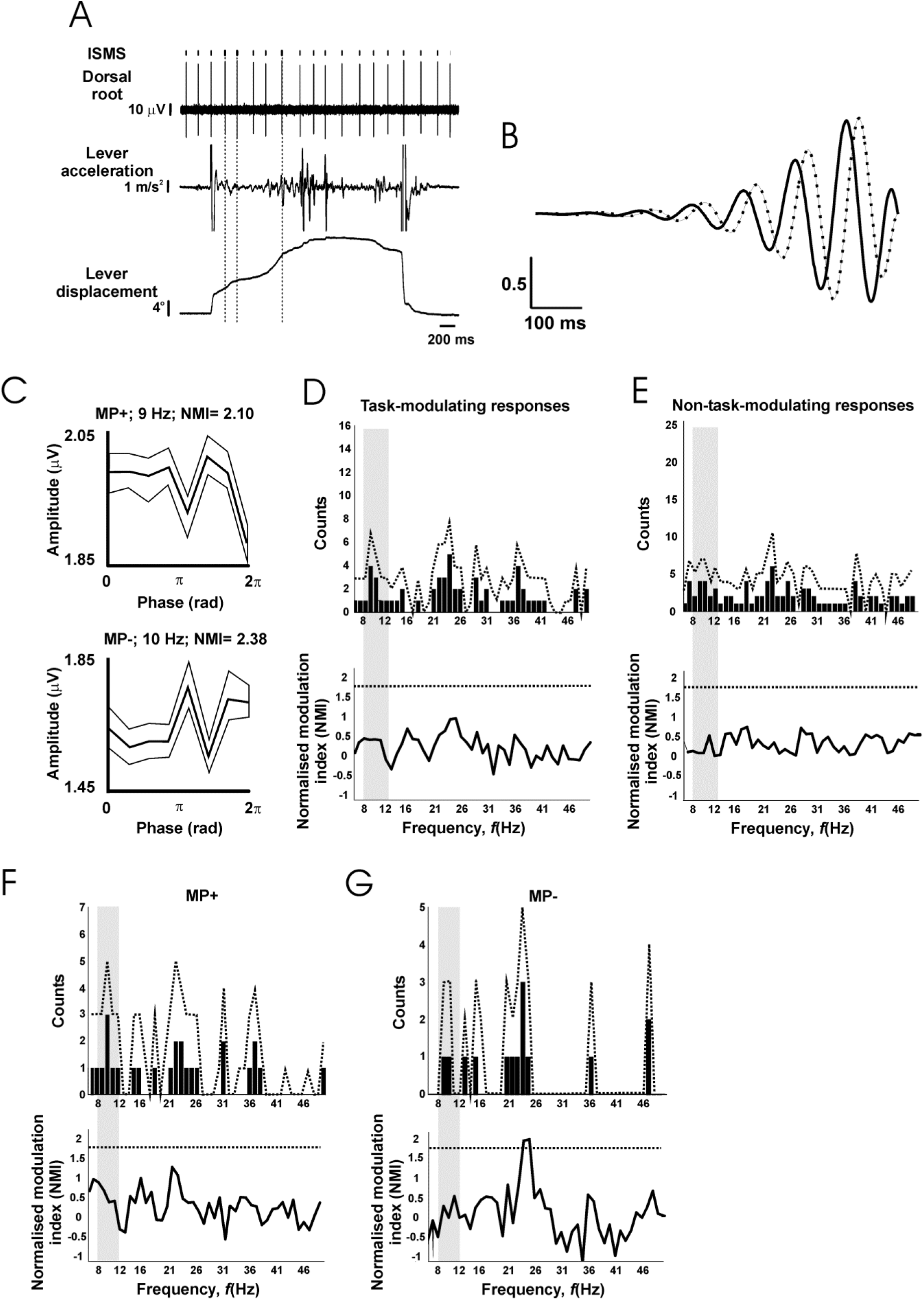
Lack of tremor-dependent modulation. A, example raw data during task performance. Vertical dotted lines indicate stimuli which passed the criteria for inclusion in analysis of tremor modulation (lever velocity>15 °/s, lever acceleration power spectral peak in the 8-12 Hz range). B, asymmetric wavelet used to determine phase of oscillations in lever acceleration; both real (solid) and imaginary (dotted) components are shown. C, example tremor modulation profiles of antidromic responses categorised on the basis of their task modulation as MP+ or MP-. Each trace shows the mean (thick line) and SEM (thin lines) of the response amplitude as a function of oscillation phase. D, number of responses which showed significant modulation with phase, as a function of frequency (top), and the mean normalised modulation index (NMI, bottom), for responses with a significant task-dependent modulation. E, as (D), but for responses without significant task-dependent modulation. F, G, as (D), but only for responses categorised as MP+ (F) or MP- (G) on the basis of their task-dependent modulation. In D-G, dotted lines indicate significance limits; traces must cross these to achieve significance (P<0.05) for an individual bin. Grey shading indicates the 8-12 Hz range relevant to peripheral tremor.

To examine whether this modulation in the ∼10 Hz tremor range was above that expected by chance and whether it reflected the specific involvement of presynaptic inhibition in regulating these frequencies, the count of significantly modulating responses and the average modulation depth across the 6-50 Hz frequency range (1 Hz resolution; see Methods) was evaluated. This is illustrated in Fig. 4D, for responses which modulated significantly with task; experimental values are shown with bars, together with the 95^th^ percentile of surrogate data with dotted traces. Neither the number of modulating responses, nor the mean modulation index exceeded the bounds expected by chance, at any of the frequencies tested. Figure 4E repeats this analysis for those responses which did not show a significant task-dependent modulation; once again, no significant modulation with tremor was detected.

In the case of the task-modulating responses of Fig. 4D, it is conceivable that pooling responses with different task-modulating profiles has obscured a significant modulation at the sub-population level. To explore this in more detail, the phase-dependent analysis was stratified by considering responses with MP+ and MP- modulating profiles separately (MP*1 and MP*2 were excluded due to their small sample size; n=2 sites each). The count of modulating sites did not rise above the bounds expected by chance for either profile at any frequency (Fig. 4FG, top traces). For the average modulation index in MP- responses, 2/45 frequency bins exceeded the 95^th^ percentile of the surrogate data, at 24 and 25 Hz. However, two or more frequency bins are expected to exceed the P<0.05 significance level merely by chance 45% of the time (surrogate distribution), therefore such modulation is not statistically significant.

## Discussion

### Physiological Mechanism Underlying Recorded Potentials

We propose that the narrow, early responses seen in the dorsal root potentials were probably generated by antidromic action potentials following stimulation of afferent axons within the spinal cord. Such responses are known to modulate if the stimulation site is close to axon terminals, because depolarisation of the terminals during pre-synaptic inhibition changes their excitability (Wall 1958). Instances where these early responses did not modulate with task could reflect either terminals which do not receive task-dependent PAD, or situations where the stimulating electrode activated stem axons, distant from the terminals and hence with a constant level of excitability. In addition, it is possible that multiple axons were activated, and that their modulation profile differed so that the modulation cancelled in the compound volley to become negligible. It is known that individual axons can show high specific patterns of pre-synaptic inhibition; even different terminals of the same axon may show different effects (Lomeli et al. 1998; Rudomin et al. 2004). However, before accepting this explanation of the likely generator of these early potentials, we must first consider their latency, which initially appears longer than expected. Measurements from photographs taken during the implant surgery of monkey V suggested a conduction distance from the point where the dorsal root leaves the cord to the first contact of the cuff electrode of 5.3 mm. Measurements from spinal sections indicated an approximate conduction distance from dorsal root to intermediate zone of 2.7 mm. Using this total conduction distance of 5.3+2.7=8.0 mm, an onset latency of 2.4 ms would imply a very slow conduction velocity of 3.3 m/s, well below accepted values for the fast cutaneous and proprioceptive afferents which are the target in these experiments. It seems unlikely that the weak ISMS (≤100 μA) delivered in these experiments could activate such slowly conducting fibres. This is supported by the behavioural reaction of the animals to these stimuli; aside from the usual brief orienting response to a stimulus, the monkeys quickly adapted and showed no signs of pain or irritation, which would be expected if we activated slow, presumed nociceptive fibres.

Several factors probably conspire to make the observed conduction longer than the naïve expectation based on conduction distance divided by expected conduction velocity. One must allow for an utilisation time of 0.1 ms, and for an additional delay due to slow conduction within the intraspinal axon terminal. For corticospinal axons, Shinoda et al. (1986) showed that the conduction velocity within terminal branches could be as low as 1 m/s. Assuming similar slowing at peripheral axon terminals, this would introduce an additional latency of around 1 ms (Baker and Lemon 1998; Shinoda et al. 1986). A further problem is that the stimulus artefact may obscure the earliest part of a response. The latencies of the visible peaks or troughs which we measure may therefore be later than the true onset latency by up to 0.5 ms (the width of an axonal action potential, Marks and Loeb (1976)). For the mean latencies observed of 2.4 ms, these considerations suggest that the conduction time in the stem axon may be only 0.8 ms, which would correspond to a velocity of 10 m/s. Finally, Loeb (1976) demonstrated that for a given axon, the conduction velocity in the dorsal root is on average 43% of that in the peripheral nerve, but this factor showed considerable variation from 20% to 70%. At the limit, a root velocity of 10 m/s would then correspond to a peripheral velocity of 50 m/s, at the upper end of the Group II range (Cheney and Preston 1976). For comparison, Seki et al. (2009) delivered ISMS and recorded volleys in the purely cutaneous superficial radial nerve peripherally; they estimated conduction velocities of 20-90 m/s, with a mean around 60 m/s. They did not correct for terminal branch and dorsal root slowing as described above in calculating these velocities, but the error is likely to have affected their readings proportionately less as their preparation had a substantially larger conduction distance (around 240 mm based on values in their Fig. 7). We conclude that the latencies of the early responses in these recordings are consistent with antidromic conduction in fast myelinated axons. It is not possible, however, to specify the nature of the axonal population; it is likely to contain a mixture of afferents responsive to both cutaneous and deep receptors.

Previous work on dorsal root potentials has used electrodes with somewhat wider spacing than that used here, and with the proximal electrode very close to the cord. Barron and Matthews (1938) reported a maximal PAD potential of 5 mV when the proximal electrode was placed on the root as it left the cord, and the distal electrode ∼10 mm away. In cat, the potential fell by half with every 1.4 mm that the proximal electrode was moved away from the cord. In this study, the cuff electrodes were located around 5.3 and 6.9 mm away from the cord. This will attenuate the recorded PAD, both because of the greater distance from the cord, and because the closer spacing will give more similar potentials which cancel in the differential recordings. Using Barron & Matthews’ estimated numbers would predict a PAD amplitude of around 200 μV. The actual recordings were substantially smaller, around 2 μV (Fig. 2B), but still detectable above the noise level by averaging. The amplitude difference presumably reflects the many differences between recordings from intact roots in awake monkey compared with roots mounted on hook electrodes in an oil pool in decerebrate cat.

In experiments in anaesthetised cats, PAD following peripheral nerve stimulation has an onset around 5-20 ms after the arrival of the volley at the cord (Eccles et al. 1962; Eccles et al. 1963b; Manjarrez et al. 2000), which corresponds to the timing of depressed synaptic transmission. In human studies reciprocal inhibition of apparent pre-synaptic origin also begins after a delay in the cord of around 5 ms (Berardelli et al. 1987). The segmental latency of the dorsal root potential recorded after afferent stimulation here was around 6 ms (Fig. 2A), which is thus compatible with PAD. By contrast, the duration of this potential was brief compared with previous reports, which often show PAD lasting tens to hundreds of milliseconds. It is possible that rather than reflecting passively conducted PAD itself, this potential reflected the antidromic discharge of axons depolarised by PAD (dorsal root reflex, Eccles et al. 1961b).

Clasically pre-synaptic inhibition is considered to reflect PAD produced by GABAergic axo-axonic synapses (Alvarez 1998; Eccles et al. 1963a). It has long been known that PAD may be also produced by extracellular potassium accumulation (Kremer and Lev-Tov 1998; Kriz et al. 1974). The terminal excitability testing used here will be sensitive to modulation in both of these mechanisms. In addition, more recent work has revealed that monoaminergic systems can induce both PAD and depression of synaptic transmission, but with no change in the excitability of intraspinal terminals to antidromic excitation (Garcia-Ramirez et al. 2014). Clearly the methods used here will fail to detect such pre-synaptic inhibition. However, monoaminergic effects on presynaptic inhibition seems to have a slow onset, with changes in synaptic transmission lagging observed changes in dorsal root potentials after 5HT application by around 20 s (Fig. 5 in Garcia-Ramirez et al. 2014). This would suggest that such mechanisms will also not be capable of temporal modulation on the timescale of tremor cycles, as we found for terminal excitability changes.

### Modulation of Pre-synaptic Inhibition

In this report, we have assumed that modulation of the antidromic volley elicited in the dorsal root by intraspinal stimulation reflects increased excitability following depolarisation of the afferent terminals, and is a marker of pre-synaptic inhibition (Wall's excitability test; Wall 1958). However, two other possibilities must also be considered. Orthodromic activity in the sensory afferents will collide antidromic spikes if the two coincide in the brief section of nerve between the spinal cord and root recording electrode. Modulation of the orthodromic firing rate with task could lead to different fractions of the antidromic spike being collided, and hence to modulation of the antidromic volley. Available data from monkeys performing a wrist flexion-extension task suggests that afferent rates modulate by around 30 discharges per second (Flament et al. 1992). Assuming a collision window equal to the antidromic response latency of the root recording (L ms), this suggests that 30 × L/1000 × 100%=3L% of antidromic spikes could collide in this way. Figure 3D presents the magnitude of the modulation of dorsal root responses following ISMS as a function of their latency; the diagonal line on that plot indicates modulation of 3L% expected if collision were the only factor involved. The majority of responses lie above this line, allowing us to conclude that collision with orthodromic spikes cannot explain all of the modulation seen.

Secondly, strong depolarisation of the afferent terminals may cause them to spike; these antidromic action potentials will render the terminal inexcitable to stimulation due to the refractory period. Such changes in excitability would still reflect changes in terminal depolarisation, although would be opposite in sign to those expected from sub-threshold depolarisation. Work in decerebrate cats walking on a treadmill reports antidromic discharge rates of around 35 Hz (Beloozerova and Rossignol 2004); assuming a refractory period of 1 ms, this would prevent responses only to 3.5% of stimuli. The modulations in Fig. 3D are generally above this level, and hence this mechanism is likely to be of little consequence for the modulation reported here.

Previous work has demonstrated that pre-synaptic inhibition of cutaneous afferents can modulate in amplitude with different phases of task performance (Seki et al. 2003; 2009), consistent with a role as a ‘gate’ to control afferent inflow during voluntary movement. The present results confirm such task-dependent modulation for a presumed mixed population of muscle and cutaneous afferents. However, based on prior work it was not clear whether pre-synaptic inhibition could modify afferent gain on a faster timescale. On the one hand, the earliest reports showed that changes in monosynaptic reflex amplitude could develop within 10 ms, and recover over around 100 ms (Eccles et al. 1961a); this work used preparations with reduced body temperature, which would plausibly have slowed the time course of effects. Under barbiturate anaesthesia at physiological temperatures there are spontaneously occurring deflections in cord-dorsum potentials. Monosynaptic reflexes evoked synchronously with these potentials are markedly potentiated, but return to baseline levels within just 30 ms (see Fig. 8 in Manjarrez et al. 2000). Fast timescale modulations in pre-synaptic inhibition therefore seem possible. On the other hand, if pre-synaptic inhibition is elicited by brief trains of stimuli, its duration can be greatly prolonged, with effects often outlasting the stimulus for up to one second (Eccles et al. 1961a; Fink et al. 2014). Even following single stimuli, effects up to 300 ms can be seen (Eccles et al. 1963b). Although the evidence is that pre-synaptic inhibition relies mainly on faster ionotropic (GABA_A_) receptors (Stuart and Redman 1992), these slower properties have been suggested to result from asynchronous release of synaptic transmitter from the axo-axonic contact, or an action on metabotropic (GABA_B_) receptors (Fink et al. 2014). Such actions would seem incompatible with fast modulation. It is not clear where within this spectrum of observations the action of physiological activity in awake behaving animals should be placed.

The present work demonstrates that, at least in one commonly occurring natural state, pre-synaptic inhibition does not modulate on fast timescales. This negative result assumes special importance in the context of previous findings related to spinal systems and their activity during the ∼10 Hz oscillations of physiological tremor. Cortical, brainstem and spinal interneuronal circuits (including pre-motoneuronal interneurons) all modulate their discharge with the tremor cycle (Williams and Baker 2009; Williams et al. 2009; Williams et al. 2010a). The phase relationship of spinal interneurons appears opposite to that of the supra-spinal centres, permitting partial cancellation of oscillatory activity at the motoneuronal level and reduction of oscillatory output. The different phase relationships appear to arise from different responses to sensory input (Kozelj and Baker 2014). Given the existence of spinal systems for phase cancelation of oscillations around 10 Hz, we must consider whether some aspect of our experimental design prevented us from detecting a modulation. The most powerful argument that this was not the case is that these results demonstrate clear modulations in spinal terminal excitability with task performance, consistent with previous work (Seki et al. 2003; 2009). Deliberately, intensities yielding responses around half-maximal were used, which should be most sensitive to modulation by excitability changes. Sufficient stimuli were delivered that the signal:noise ratio in response averages was low (Fig. 1C; see small size of error bars of Fig. 1D, 4C), arguing against statistical thresholding preventing the detection of small modulations.

One important difference between this experiment and previous work by Seki *et al.* (2003; 2009) concerns the placement of the recording site. In this work, the cuff electrode was placed on the dorsal root, meaning that recordings would contain a mixture of cutaneous and muscle afferents. By contrast, Seki *et al.* recorded from the superficial radial nerve, which has only cutaneous fibres. It is known that different categories of afferent exhibit different patterns of PAD in response to sensory or supraspinal inputs (Rudomin and Schmidt 1999). It is therefore possible that in mixed recordings different afferents modulated differently with tremor phase, leading to cancellation in the mass record and no discernible modulation. However, similar considerations would be expected to apply to task-related modulation. The fact that task-dependent effects could be seen in many recording sessions, but that tremor-related effects did not occur more than expected by chance, suggests a fundamental difference in the nature of modulation at fast versus slow timescales.

Although we found modulation of pre-synaptic inhibition during task performance, we cannot provide information on the relative contributions to this effect of afferent input versus descending control. Sensory afferents (Eccles et al. 1961a) and descending systems (Meunier and Pierrot-Deseilligny 1998; Rudomin et al. 1983) both control pre-synaptic inhibition and modulate with voluntary movement (Flament et al. 1992; Williams et al. 2010b); it is therefore likely that the observed modulations originate from changes in both afferent feedback and descending commands.

Recently, Fink et al. (2014) were able to investigate the contributions of pre-synaptic inhibition to motor control directly in mice using a genetic approach which destroyed pre-synaptic contacts, identified because they specifically express *Gad2*. During forelimb reaching, these mice show oscillatory movements which seem to result from an excessive afferent reflex gain. It would appear that pre-synaptic inhibition is modulated on relatively crude temporal timescales to mark the transition from postural stabilisation to movement, with attendant switch from a motor set dominated by reflexes to one under descending voluntary command (Seki et al. 2003; 2009; results of present work on task dependent modulation in Fig. 3). This switch allows high reflex gain during periods of constant output, but prevents reflexes from interfering with active movement. The results presented here suggest that faster modulations in afferent sensitivity in response to temporal fluctuations in output do not occur.

## Author contributions

SNB designed the study; SNB and FG carried out the experiments; FG performed data analysis and SNB and FG wrote the manuscript.

## Acknowledgements

The authors thank Paul Flecknell and Aurelie Thomas for veterinary and anaesthesia assistance, Caroline Fox and Denise Reed for theatre nursing, Terri Jackson for animal care, and Norman Charlton for mechanical workshop support.

